# NDUFV2P1-driven modulation of mitochondrial and neuronal activities; implications to schizophrenia

**DOI:** 10.64898/2026.07.23.740347

**Authors:** Yekaterina Lapiro, Rachel Karry, Ofer Binah, Dorit Ben-Shachar

**Affiliations:** Lab of Psychobiology, Department of Neuroscience, R & B. Rappaport Faculty of Medicine, Technion-Israel Institute of Technology, Israel; Department of Physiology, Biophysics and Systems Biology, R & B. Rappaport Faculty of Medicine, Technion-Israel Institute of Technology, Israel

**Keywords:** NDUFV2, NDUFV2P1, Mitochondria, Schizophrenia, Lymphocyte cell lines, Primary rat cortical neurons

## Abstract

Schizophrenia (SZ) is a severe psychiatric disorder characterized by psychosis, cognitive deficits, and disrupted social functioning. Mitochondria are essential for neuronal function, synaptic plasticity, and behavior, all impaired in SZ. A major driver of mitochondrial dysfunction in SZ is Complex I, particularly its NDUFV2 core subunit, whose alterations occur post-transcriptionally. We previously showed that the NDUFV2 pseudogene (NDUFV2P1; PG) is upregulated in brain and peripheral cells of SZ patients and inversely correlates with NDUFV2 expression and mitochondrial respiration in Epstein-Barr-virus-transformed lymphocyte cells lines (LCLs). In-silico analyses excluded small RNA interference with NDUFV2, implicating PG as an interfering factor. To study PG effects on NDUFV2 expression, mitochondrial function, and neuronal activity, we modulated PG abundance in LCLs. PG overexpression in healthy-derived LCLs impaired mitochondrial function, altering Δψm, mitochondrial network dynamics, and oxygen consumption, while PG downregulation in SZ-derived LCLs restored these parameters to normal levels. In rat cortical neurons, PG overexpression induced comparable mitochondrial disruptions alongside impaired synapse formation and reduced spontaneous neuronal firing. This study provides evidence for a mechanistic pathway through which PG interferes with NDUFV2, leading to SZ-related mitochondrial and neuronal deficits, and suggests therapeutic potential for PG downregulation in diseases with bioenergetic impairments such as SZ.

## Introduction

Schizophrenia (SZ) is a chronic psychiatric condition that typically emerges during adolescence and affects approximately 1% of the global population (1,2). Biological brain and peripheral abnormalities have been reported for this disorder (3,4). The detection of risk genes, structural brain abnormalities preceding symptom onset, subtle motor delay and cognitive deficits during the prodromal phase have contributed significantly to the conceptualization of SZ as a neurodevelopmental disorder (5–7). This disorder is characterized by abnormalities in neural circuitry, synaptic pruning and regional synaptic efficiency as well as deficits in multiple neural systems, specifically in the dopaminergic, GABAergic and glutamatergic systems (8). Notably, mitochondrial activity has also been associated with SZ. Mitochondria play a key role in energy production and calcium (Ca²⁺) buffering, both essential for proper neuronal function, supporting high energy demands and maintaining Ca²⁺ homeostasis. Studies have consistently shown a wide range of structural, biochemical, molecular, and genetic abnormalities in the mitochondria of both brain and peripheral cells in patients with SZ (9–12). Complex I (CoI)of the mitochondrial respiratory chain is considered as the rate-limiting step in overall respiration. It has been recognized as a key contributor to mitochondrial dysfunction in SZ, with its NDUFV2 subunit emerging as the most consistently and severely affected component. Notably, NDUFV2 has also been proposed as a potential risk gene for the disorder (13,14). NDUFV2 contains the N1a binuclear cluster of the N module, which functions as an antioxidant electron acceptor. NDUFV2 is a labile subunit, crucial for the proper assembly of another essential subunit, NDUFV1, which contains the NADH substrate site critical for the catalytic activity of CoI (15). It was shown that NDUFV2 loss of function is associated with increased ROS production and lipid peroxidation, decreased ATP levels in mice brains and with defective morphogenesis of hippocampal neurons dendrites and neuronal migration (16,17). Moreover, we and others have shown that a decrease in NDUFV2 levels and reduced CoI functionality led to impaired mitochondrial activity, associated with impaired neuronal differentiation, synaptogenesis and ultimately behavioral abnormalities (12,18–22). In genome-wide association studies NDUFV2 does not emerge as a strong risk gene. This observation aligns with our previous findings, which show that its DNA sequence is not changed (23), supporting the conclusion that genetic variation in NDUFV2 is unlikely to contribute to disease susceptibility. Indeed, our previous findings indicate that NDUFV2 downregulation in SZ is mediated by post-transcriptional processes. Yet, to date no experimental evidence supports the involvement of miRNAs or siRNAs in targeting NDUFV2 (24–26). Interestingly, NDUFV2 pseudogene (NDUFV2P1; PG), is significantly upregulated in SZ-derived LCLs and brain specimens. In addition, a statistically significant inverse correlation was detected between PG transcript levels and NDUFV2 transcripts and protein levels in brain and peripheral cell samples from a combined cohort of individuals diagnosed with SZ and healthy subjects. PG transcripts also showed a similar correlation with mitochondrial respiration in SZ and healthy subjects derived LCLs (23).Collectively, these data suggest that PG plays regulatory role in NDUFV2 downregulation.

Pseudogenes are DNA sequences that resemble their functional parent genes but have lost their ability to encode proteins due to mutations, deletions, or insertions. Pseudogenes are classified into three types based on their origin and the mechanisms mediating their formation, processed, unprocessed and unitary pseudogenes (27–29). The first and most common type in the human genome is processed pseudogenes, which as PG, arise from retro-transposition of their parent gene, where the mRNA is converted into DNA and inserted into a new location within the genome. Processed pseudogenes typically do not contain introns, but they possess a poly-A tail and lack regulatory sequences found in their parent genes (28,30).

Despite the inability of pseudogenes to produce proteins, many pseudogenes are transcribed into noncoding RNAs. Currently, pseudogenes, which like other noncoding RNAs, are recognized as key regulatory elements in the genome. They can influence normal cellular gene regulation and, under certain conditions, contribute to or disrupt these processes, potentially leading to pathology. Transcripts of some pseudogenes have been shown to be processed into small interfering RNAs (siRNAs) thereby silencing their parent genes (28,31), while others regulate the expression of their parent genes by competing for the same miRNAs (32–34). An additional category of pseudogenes, which share similar cis-acting sequences with their functional parent gene can compete for trans-acting molecules, primarily RNA binding proteins (RBPs) (35). Given the emerging regulatory roles of pseudogenes in health and disease, the observed increase of PG transcripts in SZ specimens, its inverse correlation with its parent gene, NDUFV2, and mitochondrial respiration, and the lack of data of other potential factor interfering with NDUFV2 transcripts (24–26) led us to investigate whether altered PG transcript levels can drive upstream abnormalities affecting NDUFV2, thereby contributing to SZ-related mitochondrial and neuronal dysfunction. If confirmed, PG could become a novel biological marker in SZ, supporting early diagnosis, elucidating disease mechanisms, and advancing the development of targeted therapeutic strategies.

## Methods and Materials

### Cells growth and maintenance

Epstein-Barr virus transformed lymphocytes cell lines (LCLs) were previously established, from control healthy subjects and DSM-IV diagnosed SZ patients (for details see Supplementary Methods) (36). LCLs were kindly provided by Prof. Bernard Lerer the Hebrew University, Jerusalem Israel and Prof. Peter Zill, Psychiatry Center, Ludwig-Maximilians-University of Munich, Germany. The study was approved by the Internal Review Board of both Centers. All patients gave written informed consent. LCLs derived from SZ- and healthy subjects either not treated (naïve) or subjected to overexpression and downregulation of NDUFV2P1 (OE PG and shPG, respectively) were studied. LCLs were randomly selected from both Israeli and Germany cohorts. Subject’s data are summarized in Supplementary Table S1A-B. All LCLs were grown as previously described (37). Typical and atypical drugs treatment of LCLs is detailed in Supplementary Methods.

Primary rat cortical neurons were isolated and grown as previously described (38). Glia cells proliferation was restrained by 4μM Arabinofuranosyl Cytidine (ARA-c, Sigma-Aldrich). All experimental protocols conformed to the guidelines of the Institutional Animal Care and Use Committee of Technion Institution and NIH.

### Protein Extraction and Immunoblotting

Protein extraction and assessment by Western blotting was performed as previously described (39,40). Antibodies are listed in Supplementary Methods.

### RNA isolation and quantification

Total RNA was isolated using TRI-Reagent (Sigma-Aldrich) and quantified as previously described (41). Total RNA was reverse transcribed into cDNA using the Verso™ cDNA kit (Thermo Scientific). The cDNA was amplified by PCR with either ReddyMix PCR Master Mix (PCR BIOSYSTENS) or FastStart Taq Polymerase kit (Sigma-Aldrich). PCR products were separated on agarose gels and quantified using the ImageQuant LAS 4000 system (GE Healthcare Life Sciences), and/or analyzed by qRT-PCR using Fast SYBR™ Green Master Mix (Thermo Scientific). All procedures followed the manufacturers’ protocols. Gene-specific primers were designed according to sequences obtained from Gene-Bank. All primers are listed in Table S2.

### Plasmids Construction, Lentiviruse production and cells infection

Human NDUFV2P1 (PG) whole transcript (NG_001161.4) was amplified from cDNA of LCLs and cloned into pEF-ENTR-A vector (Invitrogen) containing GFP. Commercially built-up shRNA plasmids (TRC2-pLKO-puro-CMV-tGFP, Sigma-Aldrich) recognizing three different sequences of PG (pLenti-shPG1/shPG2/shPG3; Table S2) were used for constructing three different PG shRNA plasmids. All generated plasmids were verified by Sanger sequencing. Using a 3rd generation packaging system (pCMV-delta-R8.2 and pCMV-VSVG plasmids), concentrated lentivirus containing each one of the above constructs was produced (Figures S1, S2) (42). The control lentivirus for OE PG, was obtained from the same plasmid lacking the PG insert (Empty), and for shPG lentiviruses a scramble sequence (Scramble). Virus titration was performed utilizing the Lenti-X GoStix Plus quantitative lentiviral titer test kit (Takara Bio), according to the manufacturer’s instructions. LCLs were infected with >1×10^5^ IFU/ml lentivirus in the presence of Polybrene (Sigma-Aldrich) and primary rat cortical neurons were infected at 7 DIV with >1×10^9^ IFU/ml lentivirus (Supplementary Methods).

### Immunofluorescence staining

Fixated and permeabilized cortical neuron cells were blocked and incubated with primary and secondary antibodies (Supplementary Methods) and then mounted with DAPI (Vectashield). Slides were viewed using Zeiss LSM 900 META Laser Scanning Inverted Confocal system (Carl Zeiss) with a x63 Plan-Apochromat oil objective. Analysis of 11–25 fields per group, obtained from 2–4 independent experiments, was performed using ImageJ software (v.1.46, Wayne Rasband). Co-localization between two fluorescent signals was defined by their spatial overlap. Pearson product-moment correlation coefficient (PCC) was assessed by the Scatterplot of co-localization analysis using ImageJ software. Branch analysis of infected neurons, identified by GFP fluorescence, was conducted using 2D/3D Skeleton analysis in ImageJ software (v1.46; Wayne Rasband), with 11 fields per group analyzed across four independent experiments.

### Oxygen consumption rates

Cellular respiration was quantified using the Seahorse Extracellular Flux Analyzer XF-96 (Seahorse Biosciences) as previously described (43). Oxygen consumption rates (OCR) were measured before and after the addition of various inhibitors to determine key parameters of mitochondrial respiration (Supplementary Methods; Figure S3) (44,45). Optimized concentrations of respiration inhibitors were: 4 µM oligomycin, 1 μM FCCP and 3 μM Rot/AA for LCLs and 2 µM oligomycin, 1 μM FCCP and 3 μM Rot/AA for primary rat cortical neurons. Measurements were made in 8-12 replications of n=5 LCLs/group. Data were normalized to cells’ protein levels in each well of the Seahorse plate using the modified Lowry method (46).

### Mitochondrial imaging

Mitochondrial membrane potential (Δψm), their cellular distribution and network connectivity analyses were performed using MitoTracker® Orange CMTMRos mitochondrion-selective probes (Invitrogen) as previously described (29,43) and detailed in Supplementary Methods. Images were acquired using Zeiss LSM-700 or LSM-900 Laser Scanning Confocal System (Carl Zeiss Microimaging) with a x63 Plan-Apochromat oil objective and analyzed by in-house Python scripts. 3D visualization and data analysis were also performed using IMARIS software. Data were obtained from 6-28 cells/group in n=2 LCLs/group in at least two independent experiments.

### Microelectrode array (MEA)

For network activity recording, 5×10^5^ cells were seeded on Laminin/poly-D-lysine (Sigma-Aldrich) covered thin glass multielectrode array (MEA) dishes, grown and infected at 7 DIV. Standard MEA dishes containing 60 electrodes (30 μm titanium nitride electrode diameter) in an 8 by 8 grid arrangement spaced 200 μm apart with a small glass ring, were used. Three to four cultures plates/group obtained from 2 independent experiments were recorded at 15-22 DIV, minimum 8 days after infection. Recording of the spontaneous activity was performed for 10 min with a sampling frequency of 20 kHz. Data of individual electrodes were collected using the MC_Rack software (version 1.21 by Multi Channel Systems MCS GmbH). Field potentials of raw data were corrected with a high pass filter of a frequency of 200Hz. The spontaneous activity was detected by a spike detector algorithm consisting of a hard threshold crossing, computed using four times the standard deviation of the raw signal. The bursts were defined as sequences of three spikes occurring in less than 100 msec. Electrophysiological data were imported to MATLAB software (MathWorks) and analyzed using an in-house script. (47–50)

### Statistical analysis

Results are expressed as mean ± SEM. A power analysis was performed to estimate the required sample size. Data were analyzed for normal distribution by Kolmogorov-Smirnov test. Normally distributed data were analyzed by unpaired or paired Student T-test. Differences in means between groups were considered significant if p≤0.05. Samples distance of at least ±2 SD from group average were excluded from the analysis. All Statistical analyses, except for LCLs confocal mitochondrial imaging, were performed using GraphPad Prism 9.1.1. For the analysis of LCLs confocal mitochondrial imaging, mixed-effects model tests were performed, and effect sizes (Cohen’s d) with confidence intervals (CI) were calculated using Python.

## Results

Our previous studies suggest that the increased levels of PG transcripts in SZ specimens underlies the deficits observed in its parent gene, NDUFV2. To examine the effects of changes in PG/NDUFV2 expression ratio on mitochondrial function, we overexpressed PG (OE PG) in healthy cells to mimic PG state in naïve SZ cells and downregulated PG (shPG) in SZ-LCLs, to restore a healthy cellular state. Infection efficiency was >85% as determined by GFP fluorescence intensity. (Figure S5). First, we verified that our selected LCLs cohorts show a significant decrease in the mRNA and protein expression of NDUFV2 and an increase in mRNA expression of PG in SZ-derived LCLs compared to healthy LCLs (P<0.02, P<0.005 and P<0.003, respectively; Figure 1A, B, G, J). Our preliminary results suggest that this pattern of change is not due to typical (haloperidol) or atypical (clozapine and risperidone) antipsychotic drug treatment, as 14 days of *in-vitro* exposure of LCLs to these drugs did not affect the transcript levels of either the pseudogene or NDUFV2 (Figure S6).

**Figure 1:**
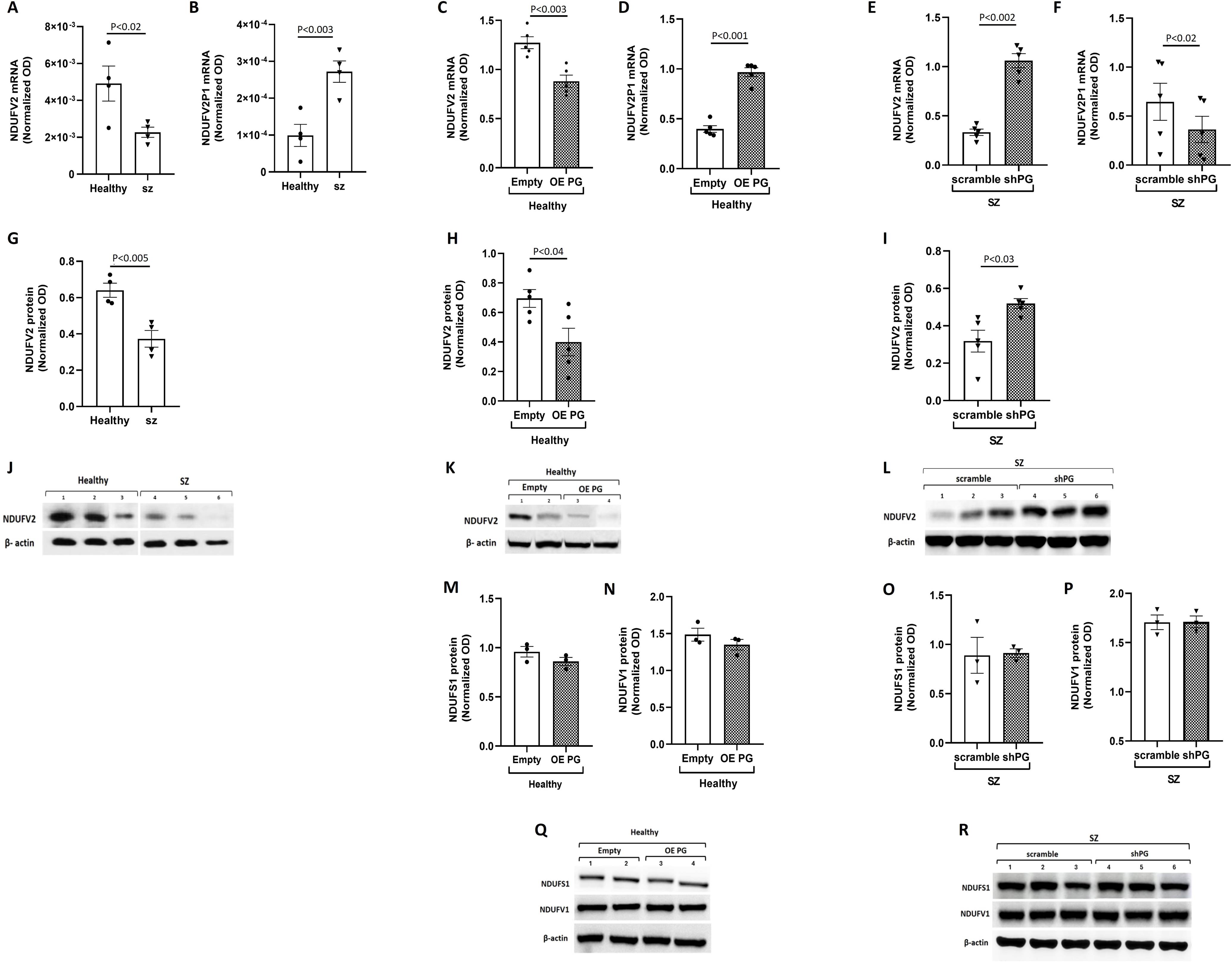
NDUFV2P1 (PG) affects its parent gene NDUFV2 transcript and protein levels. Naïve SZ-derived LCLs showed a significant decrease (P<0.02, n=4 LCLs/group) in NDUFV2 **(A),** while an increase (P<0.003, n=4 LCLs/group) in PG mRNA levels as compared to naïve healthy subjects-derived LCLs **(B)**. Overexpression of PG (OE PG) in healthy subject-derived LCLs caused a significant decrease (P<0.003, n=5 LCLs/group) in NDUFV2 **(C)** and a significant increase in PG (P<0.001, n=5 LCLs/group) transcript levels as compared to the same LCLs infected with an empty virus (Empty) **(D)**. Downregulation of PG (shPG) in SZ-derived LCLs, caused a significant increase (P<0.002, n=5 LCLs/group) in NDUFV2 mRNA levels **(E)** and a significant decrease (P<0.02, n=5 LCLs/group) in PG as compared to the cells infected with scramble virus **(F)**. Representative gels and quantification of the results demonstrated a significant reduction (P<0.005, n=4 LCLs/group) in NDUFV2 protein levels in naïve SZ-derived LCLs as compared to healthy subjects -derived LCLs **(G, J)**. OE PG in healthy subjects-derived LCLs initiated a significant decrease (P<0.04, n=5 LCLs/group) in NDUFV2 protein levels **(H, K)**, while shPG treatment in SZ-derived LCLs, caused a significant increase (P<0.03, n=5 LCLs/group) in NDUFV2 protein levels as compared to the cells infected with scramble virus **(I, L)**. Overexpression **(M-N**, **Q)** or downregulation **(O-P**, **R)** of PG in healthy and SZ- derived, respectively, had no effect on protein levels of NDUFV1 and NDUFS1 (n=3 LCLs/group), two additional proteins of the N module of CoI. Values are expressed as mean ± SEM normalized to β-actin. Data was obtained from n=3-5 LCLs/group in two independent experiments.

OE PG in healthy-derived LCLs resulted in a significant reduction of NDUFV2 mRNA and protein levels (P<0.003 and P<0.04, respectively; Figure 1C, H, K). These reductions were associated with a significant increase in PG transcript levels following OE PG-Lenti treatment compared to cells infected with Empty virus (P<0.001; Figure 1D) mimicking the findings in naïve SZ-derived LCLs. However, shPG treatment in SZ-derived LCLs resulted in a significant increase in NDUFV2 mRNA and protein levels (P<0.002 and P<0.03, respectively; Figure 1E, I, L), associated with a significant reduction in PG transcript levels (P<0.02; Figure 1F). Both OE PG and shPG had no effect on NDUFS1 and NDUFV1 subunits’ protein levels (Figure 1M-R), additional subunits of CoI, which together with NDUFV2 form the functional N module of CoI (51,52) and have been reported to be altered in SZ (12).

Naïve SZ-derived LCLs demonstrated deficits in mitochondrial function. Hence, we observed a significant reduction in mitochondrial respiration parameters including basal-, ATP-linked and maximal- cellular oxygen consumption rates (OCR) (P<0.02, P<0.03 and P<0.05, respectively; Figure 2A-C). This decrease in OCR parameters was also manifested at the level of mitochondrial membrane potential (Δψm) (P<0.001, Cohen’s d=2.97, 95% CI [2.47, 3.69]; Figure 2E, F).

**Figure 2:**
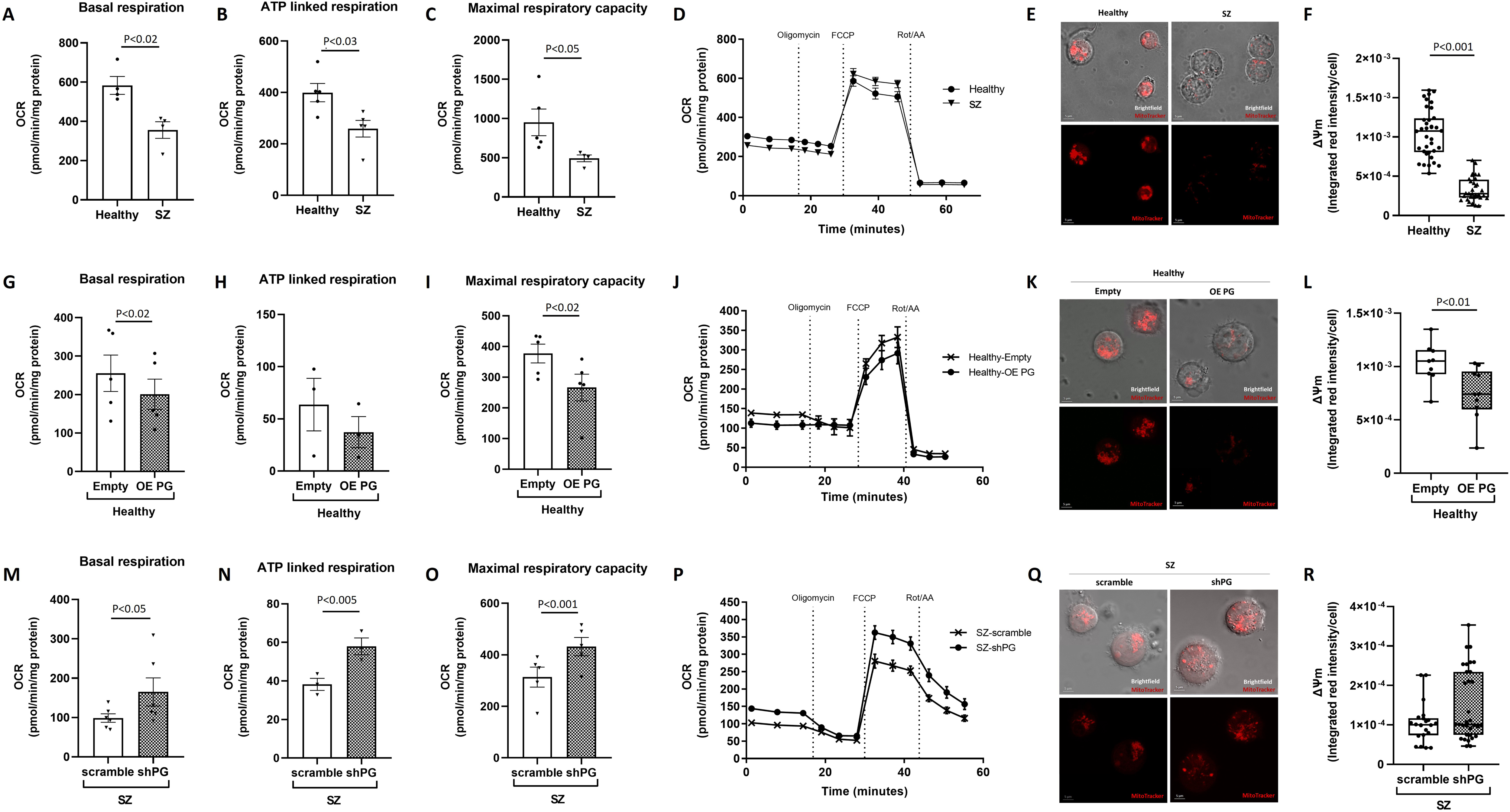
Alterations in NDFV2P1 (PG) transcript levels impact mitochondrial functions. Quantifications of oxygen consumption rates (OCR) in naïve SZ and healthy control LCLs (**A-D**), following OE PG and its Empty control in healthy-derived LCLs (**G-J**) and following shPG and its scramble control in SZ-derived LCLs (**M-P**). Naïve SZ cells exhibited significantly reduced levels of all tested respiratory parameters including the basal, ATP-linked and maximal OCR as compared to healthy LCLs (P<0.02, P<0.03 and P<0.05, respectively) **(A-C)**. As a result of OE PG in healthy LCLs, a significant decrease was observed in basal and maximal OCR (P<0.02 and P<0.02, respectively) **(G, I)**, while no significant change was observed in ATP-linked OCR **(H)**. Following shPG in SZ-derived LCLs, a significant increase in basal, ATP-linked, and maximal OCR was observed (P<0.05, P<0.005 and P<0.001, respectively) **(M-O**). OCR profiles following the addition of oligomycin, FCCP and Rot/AA with and without manipulation of PG expression **(D, J, P)**. OCR measurements were done in 8-12 replications/LCL n=5 LCLs/group. All values are expressed as mean ± SEM. Representative images of MitoTracker Orange fluorescence concentrated in mitochondria depending on their membrane potential (Δψm) within LCLs (**E, K, Q**). Quantifications of Δψm in healthy control and naïve SZ LCLs **(F)**; following PG overexpression (OE PG) and its Empty control virus (Empty) in healthy-derived LCLs (**L**); and following PG downregulation (shPG) and its scramble control virus (scramble) in SZ-derived LCLs (**R**). Δψm is significantly decreased (P<0.001, Cohen’s d=2.97, 95% CI [2.47, 3.69]) in naïve SZ cells and in healthy-derived LCLs following OE PG (P<0.01, Cohen’s d=1.29, 95% CI [0.30, 2.28]), while no change was observed in Δψm following shPG in SZ-derived LCLs, compared to their relevant controls. Data were analyzed using a linear mixed-effects model, with cells (n=8-24 per line) treated as repeated measures nested within each line (n=2 LCLs/group), and line modeled as a random effect.

We then investigated whether PG- induced alterations in NDUFV2 projects on mitochondrial function. Indeed, OE PG caused a significant decrease in basal and maximal OCR (P<0.02 and P<0.02, respectively; Figure 2G, I) as well as in Δψm (P<0.01, Cohen’s d=1.29, 95% CI [0.30, 2.28]; Figure 2K, L) in healthy-derived LCLs as compared to Empty infected cells. No significant change was observed in ATP-Linked OCR (Figure 2H). The functional link between PG and mitochondrial activity was further supported by shPG treatment in SZ-derived LCLs. A significant increase was observed in basal, ATP-linked, and maximal OCR (P<0.05; P<0.005 and P<0.001, respectively; Figure 2M-O). While Δψm remained unchanged following shPG treatment (Figure 2Q, R). Figure 2D, J, P represents the OCR profile following the addition of oligomycin, FCCP and Rot/AA with and without manipulation of PG expression.

Mitochondrial connectivity and distribution within the cell are crucial for their function. These organelles dynamically undergo fusion and fission in response to cellular demands, playing a vital role in mtDNA maintenance, metabolic energy regulation, stress response signaling, and spatial organization. (53,54). In naïve SZ LCLs mitochondria network connectivity was significantly decreased (P<0.05, Cohen’s d=1.11, 95%, CI [0.62, 1.6]); Figure 3A, B) and the mitochondria were unevenly distributed throughout the cells, indicated by higher coefficient variation (P<0.05, Cohen’s d=0.97, 95% CI [0.43, 1.51]); Figure 3C, D), as compared to healthy-derived LCLs. OE PG in healthy cells caused a similar phenomenon in network connectivity (P<0.05, Cohen’s d=0.95, 95% CI [0.046, 1.85]); Figure 3E, F) but had no effect on mitochondrial cellular distribution (Figure 3G, H). In shPG infected SZ LCLs however, we observed a significant increase in the mitochondrial network connectivity (P<0.01, Cohen’s d=1.3, 95%, CI [0.27, 2.32]); Figure 3I, J) as well as improvement in cellular mitochondrial distribution, indicated by a decrease in the coefficient of variation, (P<0.04, Cohen’s d=0.97, 95% CI [0.24, 1.71]; Figure 3K, L) as compared to SZ-LCLs infected with the control vector. Consistent with the network connectivity data, MFN2, OPA1 and DRP1, three key players in mitochondrial fusion and fission, were significantly reduced following overexpression of PG in healthy LCLs (P<0.05, P<0.004 and P<0.03, respectively; Figure 3M-P), while significantly increased upon downregulation of PG in SZ LCLs (P<0.0001, P<0.03 and P<0.01, respectively; Figure 3Q-T).

**Figure 3:**
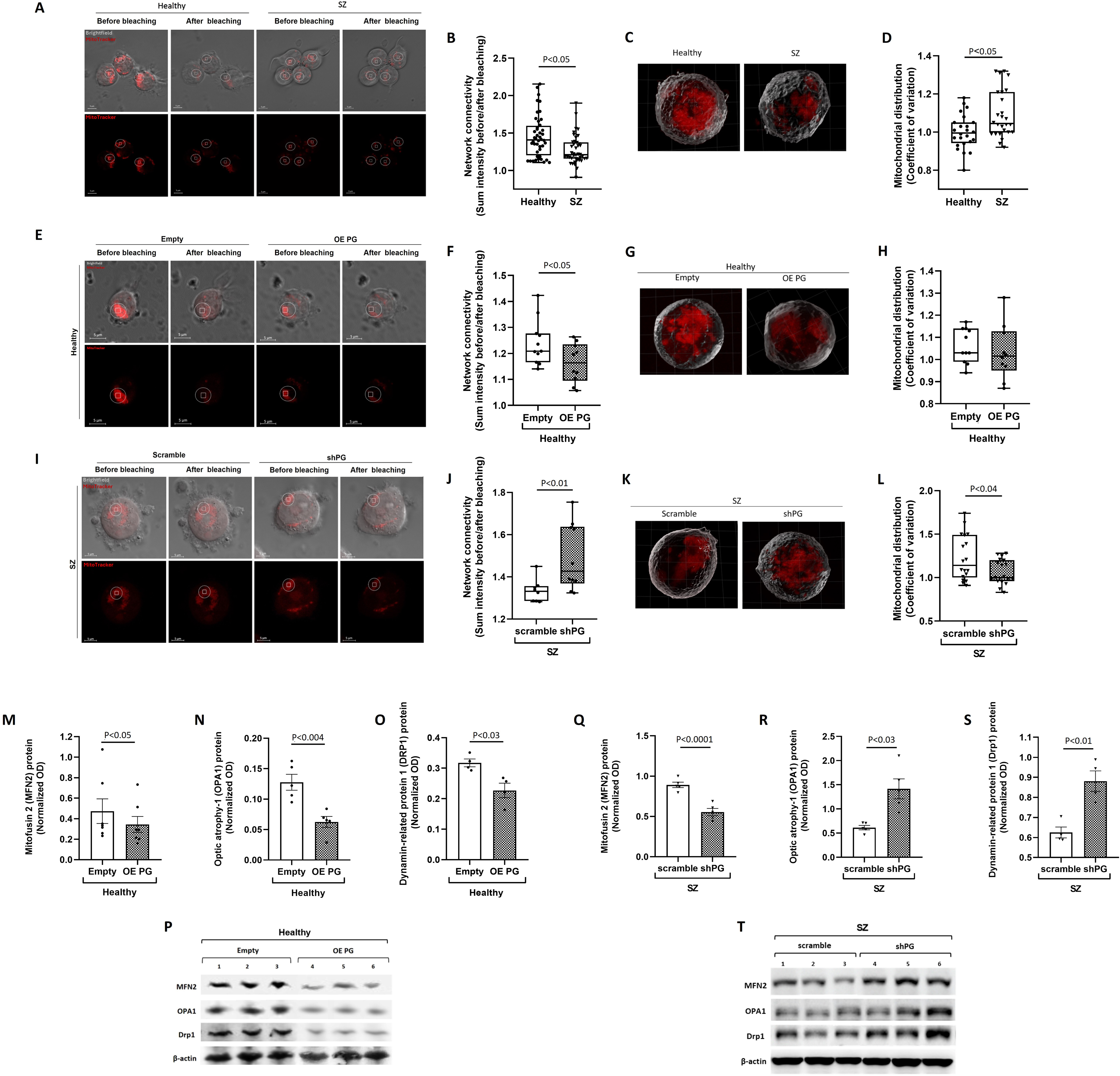
Alterations in NDUFV2P1 transcript levels affect mitochondrial network connectivity, distribution and fission/fusion protein expressions. Representative images of mitochondria stained with Orange MitoTracker before and after focused laser beam-induced dissipation of Δψm. The square marks the area treated with the focused laser beam. The dashed circle (ROI) marks the area within and beneath which the 3D fluorescence intensity before and after laser beam application was quantified in naïve SZ and healthy LCLs **(A)**; following OE PG and its Empty control in healthy-derived LCLs (**E**); and following shPG and its scramble control in SZ-derived LCLs **(I)**. Mitochondria network connectivity was significantly reduced in naïve SZ-derived LCLs compared to healthy subjects (P<0.05, Cohen’s d=1.11, 95%, CI [0.62, 1.6]) **(B).** OE PG in healthy-derived LCLs caused a significant reduction in mitochondria network connectivity as compared to Empty infected cells (P<0.05, Cohen’s d=0.95, 95% CI [0.046, 1.85]) **(F)**, Conversely, a significant intensification of mitochondrial connectivity was observed following shPG treatment in SZ-derived LCLs as compared to scramble infected cells (P<0.01, Cohen’s d=1.3, 95%, CI [0.27, 2.32]) **(J)**. Representative 3D images obtained by IMARIS software, of MitoTracker Orange stained mitochondria in naïve SZ and healthy LCLs, and in OE PG and shPG treated healthy and SZ derived cells, respectively **(C, G, K)**. Mitochondrial distribution within the cell borders was determined by coefficient of variation between the grid intensity of the pixels. The coefficient of variation was elevated in naïve SZ-derived LCLs compared to healthy subjects (P<0.05, Cohen’s d=0.97, 95% CI [0.43, 1.51]) **(D)**, while no change was observed in OE PG healthy-derived LCLs **(H)**. shPG in SZ-derived LCLs caused a significant decrease in the coefficient of variation (P<0.04, Cohen’s d=0.97, 95% CI [0.24, 1.71]) **(L)**. All confocal data were analyzed using a linear mixed-effects model, with cells (n=6-28 per line) treated as repeated measures nested within each line (n=2-3 LCLs/group), and line modeled as a random effect. Following OE PG infection in healthy-derived LCLs, levels of the fission/fusion proteins MFN2, OPA1, and Drp1 were significantly decreased (P<0.05, P<0.004, and P<0.03, respectively) **(M-O)**. Conversely, their levels were significantly increased (P<0.0001, P<0.03, and P<0.01, respectively) following shPG infection of SZ-derived LCLs **(Q-S)**. Representative SDS-PAGE gels of fission/fusion protein following PG OE and shPG **(P, T)**. Values are expressed as mean ± SEM following normalization to β-actin of n=4-5 LCLs/group of two independent experiments.

To assess the effects of PG in neurons, we overexpressed the human PG (hPG) in primary rat cortical neurons as in-silico analysis revealed that NDUFV2 has no pseudogene in rats. Following hPG overexpression (OE PG) (P<0.02; Figure 4A), we observed a significant reduction in the NDUFV2 protein levels (P<0.001) yet not in the two additional subunits of CoI, NDUFS1 and NDUFV1, similar to our finding in LCLs. The transcript level of NDUFV2 was significantly increased (P<0.001), which most likely stems from compensatory mechanisms addressing neuronal cells’ high energy demands (55,56); (Figure 4B-F). Concomitant with our findings in LCLs, OE PG caused alterations in mitochondrial function. Hence, Δψm was significantly decreased and the mitochondria were unevenly distributed as indicated by the increase in the coefficient of variation (P<0.03 and P<0.001, respectively; Figure 4G-I). No change was observed in mitochondria network connectivity in cells’ soma following OE PG (Figure 4J), similar to our previous finding in SZ-iPSCs-derived glutamatergic neurons (19). Basal, ATP-linked and maximal OCR were also significantly reduced (P<0.0001, P<0.0001 and P<0.0004, respectively; Figure 4K-N). Notably, the effects of OE PG did not involve ROS formation, as the treatment of the infected cultures with NAC did not change the basal OCR levels in OE PG, unlike the reduction observed upon culture treatment with H_2_O_2_ (Figure S7).

**Figure 4:**
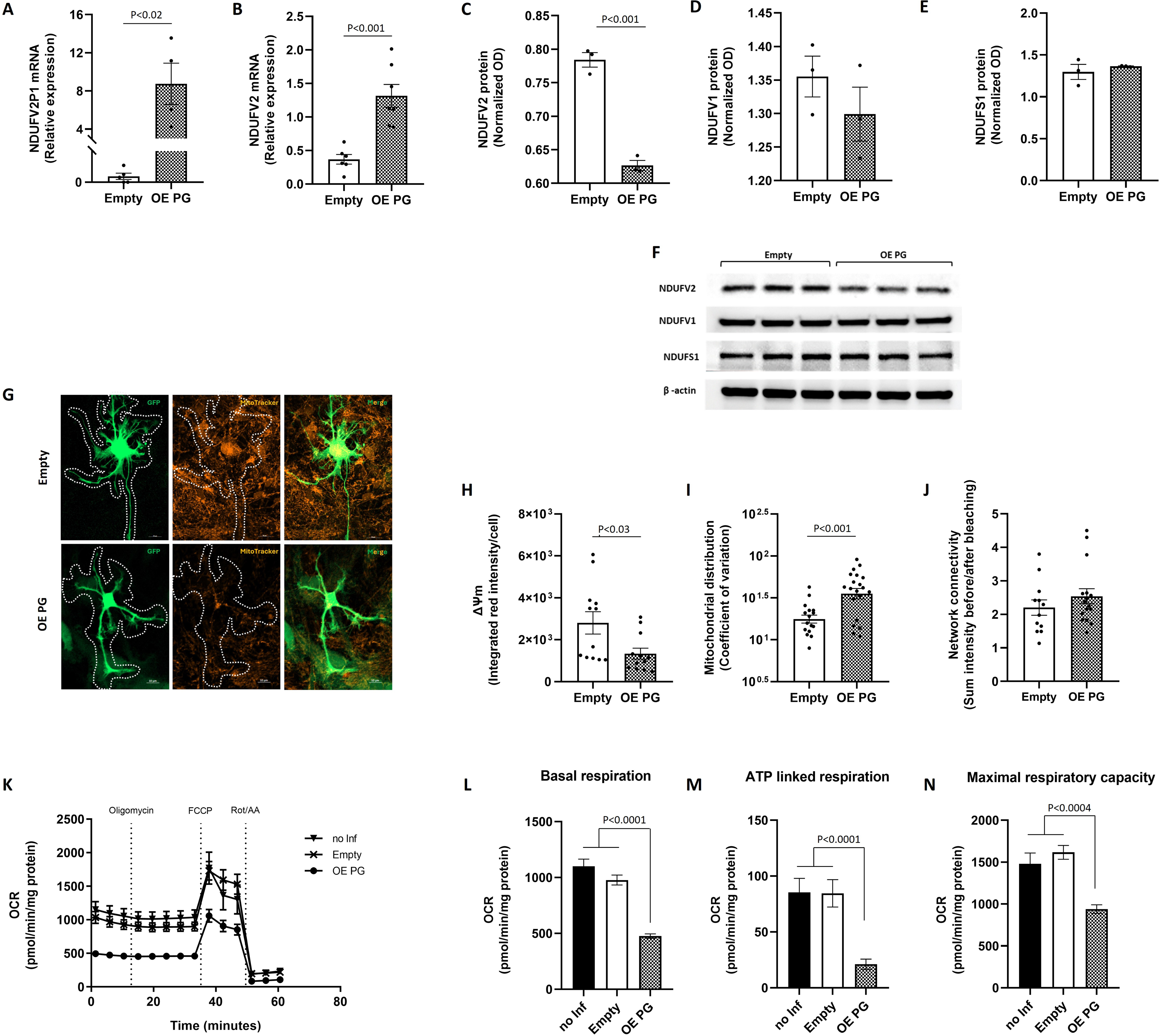
Overexpression of human NDUFV2P1 (OE PG) in primary rat cortical neuronal cultures impacts its parent gene, NDUFV2, and mitochondrial functions. Quantification of both hPG and rat NDUFV2 transcript levels exhibited a significant increase following OE PG (P<0.02 and P<0.001, respectively) **(A-B)** while a significant decrease (P<0.001) was observed in protein levels of NDUFV2 with no changes in NDUFV1 and NDUFS1, as represented by SDS-PAGE gels and their quantifications **(C-F)**. Representative images of MitoTracker Orange fluorescence, representing Δψm within infected primary cortical neurons marked with GFP (green) acquired using Zeiss LSM 900 Laser Scanning Confocal System **(G)**. Quantification of Δψm in the selected ROI (indicated by the dashed line) revealed a significant decrease (P<0.03) following OE PG compared to Empty infected cells **(H)**. The distribution, determined by coefficient of variation between the grid intensity of the pixels within the ROI, was significantly elevated (P<0.001) following OE PG **(I)**. No change was observed in mitochondria network connectivity **(J)**. OCR profiles following OE PG, Empty infected and non-infected (no Inf) cells in the presence and absence of oligomycin, FCCP, and Rot/AA are presented in **(K)**. OE PG caused a significant impairment in all analyzed cellular respiration parameters compared to both Empty infected and no Inf cells, in basal, ATP-linked and Maximal OCR (P<0.0001, P<0.0001 and P<0.0004, respectively) **(L-N)**. Values are expressed as mean ± SEM of 8-16 replications/group in 2 independent experiments. For analysis of β-actin normalized transcript and protein, values are expressed as mean ± SEM of n=3 or 6/group, performed in 2 independent experiments. For imaging analysis values are expressed as mean ± SEM of 12-13 cells/group, performed in two independent experiments.

We further studied whether OE PG-induced mitochondrial dysfunction leads to neuronal aberrations. OE PG in rat primary cortical neuronal cultures caused structural changes in neurons as represented by the binary images of neurons, and by the significant decrease in both neuronal number of branches (P<0.004) and junctions (P<0.004) (Figure 5A-C). No change was observed in the average length of the branches (Figure 5D). Quantification of the presynaptic and postsynaptic markers, SYN1 and PSD95 respectively, showed a significant decrease expressed by puncta density (P<0.03 and P<0.001, respectively; Figure 5E-G) of both markers. The co-localization of the pre- and postsynaptic markers, indicating synaptic contacts, was also significantly reduced (P<0.0001; Figure 5H). This was substantiated by Pearson product-moment correlation coefficient (PCC) showing a clear difference in signal overlap between Empty and OE PG neurons, with corresponding r values indicating reduced co-localization upon OE PG treatment (r=0.0305 in OE PG versus r=0.659 in Empty; Figure 5I). Finally, these neuronal changes were complemented by a significant decrease in neuronal basal firing rates, manifested by the reduced spike and burst frequencies following treatment with OE PG (P<0.01 and P<0.001, respectively; Figure 5J-L).

**Figure 5:**
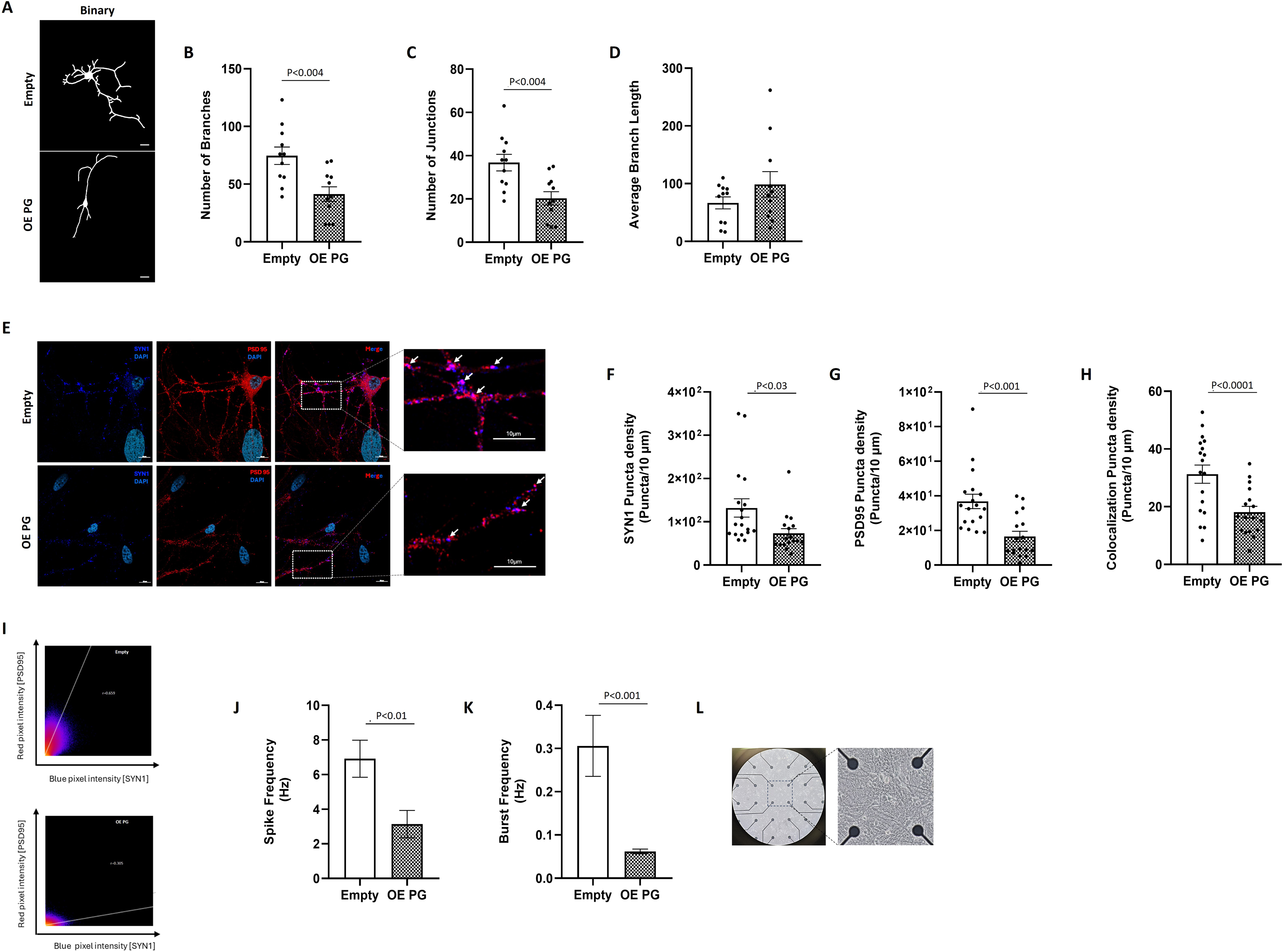
Overexpression of hPG in primary rat cortical neuron cultures impairs neuronal sprouting, synapse formation, and basal firing activity. Representative binary images of neurons following Empty and OE PG infection **(A)**. The qualifications of images reveal a significant decrease (P<0.004) in both the number of branches and junctions **(B-C)** while no change was observed in the average length of the neuronal branches **(D)**. Results were obtained from 11 fields per group across four independent experiments and analyzed using 2D/3D Skeleton analysis in ImageJ software. Representative Confocal images of the pre-synaptic marker SYN1 (blue) and post-synaptic marker PSD-95 (red) and their merged image following OE PG and Empty infection of 14 DIV primary cortical neuron cultures. The images on the right are a magnified view of the area outlined by the dashed white line in the adjacent image. Each arrow indicates co-localization of the two markers SYN1 and PSD95 **(E)**. Quantification of SYN1 and PSD95 puncta densities revealed a significant reduction following OE PG infection (P<0.03 and P<0.001, respectively), and a marked decrease in their co-localized puncta density (P<0.0001) **(F-H)**. A representative image of Pearson product-moment correlation coefficient (PCC) demonstrates a marked reduction in signal overlap between SYN1 and PSD95 following OE PG treatment compared to the Empty control (r=0.0305 vs. r=0.659) **(I)**. Values are expressed as mean ± SEM of 11-18 cells/ group and performed in two independent experiments. Recordings of the spontaneous activity of neurons were collected using the sofware MC_Rack. A significant decrease in spikes **(J)** and bursts **(K)** frequencies was observed (P<0.01 and P<0.001, respectively) after treatment with OE PG compared to Empty control. Representative image of neuronal network in the Microelectrode Array (MEA) plate is presented **(L)**. Values are expressed as mean ± SEM of 3-4 plates/ group, performed in two independent experiments.

## Discussion

Increased NDUFV2P1 (PG), while decreased NDUFV2 expression was previously demonstrated in brain specimens and lymphocytes of patients with SZ (23). Although a highly significant negative correlation was observed between the two, the link between both and the upstream effects due to their aberrant cellular ratio is not clear. The findings of the present study show that PG negatively regulates the expression of the NDUFV2, thereby leading to mitochondrial dysfunction, disrupted neuronal structure and reduced spontaneous firing, all impaired in SZ.

The full spectrum of changes induced by PG was revealed by its overexpression (OE PG) and downregulation (shPG) in healthy and SZ-derived LCLs, respectively, both disease-relevant states. By overexpressing PG in healthy-derived LCLs, we were able to simulate the mitochondrial-related abnormalities observed in naïve SZ-derived LCLs. Conversely, downregulation of PG (shPG) restored mitochondrial deficits observed in SZ-derived LCLs. Notably, the manipulation of PG levels specifically affected the NDUFV2 subunit of CoI N functional module with no change in its two other subunits, NDUFV1 and NDUFS1. The lack of change in NDUFS1 is in line with our previous studies in SZ human brain, peripheral cells and LCLs total lysate. NDUFV1, however, was altered in brain and peripheral samples of SZ patients but not in SZ LCLs similar to the results of this study (23,57,58). These data substantiate the specificity of PG effect on the N module of CoI.

The imbalance in cellular PG/NDUFV2 ratio by overexpression or downregulation of PG, affected major mitochondrial function as demonstrated by altered cellular oxygen consumption rates (OCR), and mitochondrial network connectivity. Mitochondrial function is interconnected with its network dynamics (59). Hence, it was previously reported that loss of CoI activity and impaired NAD⁺ regeneration, both important factors of the electron transfer chain, were associated with reduced levels of the fusion protein OPA1 (60,61). In addition, in rats exposed to hyperoxia, which reduced ATP production, an increase in mitochondrial fission via the DRP1 signaling pathway was observed (62). On the other hand, disruption of the fusion and fission proteins MFN2, OPA1 and DRP1 led to mitochondrial heterogeneity and reduced cellular respiration (63). Concomitantly, PG-induced mitochondrial dysfunction was associated with reduced levels of OPA1, MFN2, and DRP1 following OE PG in healthy-derived LCLs, while their increase following shPG-induced restoration of mitochondrial function in SZ-derived LCLs. The interaction between mitochondria and fission/fusion proteins suggests an activation of a vicious/virtuous circle that can amplify the PG-induced negative/positive effects on mitochondrial function. Mitochondrial Δψm, the driving force for ATP production, was altered only following PG overexpression (OE PG) in healthy LCLs but not after its downregulation (shPG) in SZ-LCLs. This counterintuitive finding, given the enhanced OCR, calls for further research focusing on the coupling mechanisms between oxidation and phosphorylation in shPG infected cells.

We have also overexpressed PG in SZ-derived LCLs and downregulated it in healthy-derived LCLs. OE PG in SZ cells worsens almost all the studied mitochondria related parameters. However, silencing of PG in healthy-derived LCLs using the same experimental conditions as in SZ-derived LCLs failed, probably due to their initial low levels of PG. Nevertheless, the overexpression of PG in healthy subjects-derived LCLs and its downregulation in SZ-derived LCLs substantiate our main hypothesis of PG involvement in mitochondrial functional outcomes.

The experimental model used in this study is LCLs derived from patients and healthy subjects. Although LCLs may not replicate the full complexities of tissue environment in a patient, they are conceptualized as retaining the genetic and epigenetic characteristics relevant to the disease. SZ exhibits peripheral manifestations including altered membrane phospholipids and impaired amino acid transport in fibroblasts and muscle and mitochondrial dysfunction in blood cells (57,64,65). Consequently, LCLs have been suggested as a medication-free peripheral model for neurological and psychiatric disorders (69,70), turning LCLs into a reliable and valuable model for studying biological abnormalities in SZ.

Next, we investigated whether the effect of PG on mitochondria projects onto neuronal function, given the substantial evidence supporting a close link between mitochondria and neuronal activity and connectivity (71–73). For example, mitochondrial dysfunction was associated with deficits in neuronal sprouting, synaptic connectivity, and neurotransmission throughout the differentiation of dopaminergic and glutamatergic neurons (20). Furthermore, a recent elegant study showed that mitochondrial energy metabolism efficiency significantly affected the developmental tempo of cortical neurons (74). Hence, enhancing mitochondrial metabolism in human neurons by transplanting metabolically more efficient mouse mitochondria accelerated their maturation, enabling cells to reach an advanced developmental stage earlier than usual and vice versa. Mitochondrial impact on neuronal function, and thereby behavioral responses, is further supported by previous transplantation studies of normal active mitochondria in SZ, Alzheimer’s and Parkinson’s disease rodent models (75–77).

Here, we demonstrate that PG is able to modify neuronal parameters, which were also affected by mitochondrial transplantation (19). Thus, overexpression of human PG (OE PG) in rat primary cortical neurons, a core brain region implicated in SZ, showed deficits in all studied mitochondrial parameters similar to those observed in OE PG healthy-derived LCLs. These changes were probably not due to a state of oxidative stress induced by OE PG, as adding NAC did not improve the OCR parameters. Cortical neurons that overexpressed PG and showed mitochondrial deficits demonstrated structural abnormalities expressed by a significant reduction in the number of branches and junctions. Previous reports suggest that the spatial morphology of neurons alone can provide insights into the nature of their activity (78). Indeed, these neurons presented a reduction in both the pre- and post-synaptic markers, SYN1 and PSD95, respectively, and their co-localization, an indicator of synapse formation. These findings are in line with previous studies that reported an association between a reduced number of mitochondria in dendrites and a decrease in the number of synapses and dendritic spines. On the other hand, boosting mitochondrial content or activity led to an increase in spines’ number and synapse plasticity (79,80). Finally, the basal activity of the neuronal network in which PG was overexpressed was significantly decreased as manifested by the reduction in their spikes and burst frequencies. Given that no intrinsic pseudogene of NDUFV2 has been identified in rats, the suppression of rat NDUFV2 translation associated with OE PG, which further projects to mitochondrial and neuronal functions, is likely attributed to the high homology (65%) between their mRNAs. In line with the latter, our unpublished data show that PG is a trans-acting molecule that interferes with NDUFV2 transcripts on RNA binding proteins.

In all, the data of this study demonstrate that an imbalance in cellular PG/NDUFV2 ratios serves as a bottom-up driving force for mitochondrial impairment, which in turn disrupts neuronal synapse formation and firing, pathologies observed in SZ. The question whether PG/NDUFV2 imbalance affects behavior, and the mechanisms through which PG interact with NDUFV2, are currently under investigation and may reveal additional therapeutic targets in SZ.

Whether the reduced level of NDUFV2 following upregulation of NDUFV2P1 is a cause or a consequence of SZ symptomatology remains unclear. We suggest that diverse abnormalities observed in SZ such as synaptic, neuronal, and immune dysfunction, are interwoven and together contribute to the disease symptoms. Indeed, we have previously shown that the beneficial/toxic molecular and behavioral outcomes of mitochondrial transplantation depend on the physiological state of the recipient (e.g. immune, energy metabolism, hormonal states). Thus, transplantation of healthy mitochondria into the medial prefrontal cortex of a rodent model of SZ improved behavioral symptoms, while eliciting adverse effects in healthy controls (75). The potential toxicity of mitochondria has reduced the likelihood of their transplantation as a therapeutic strategy for SZ, and instead supports the consideration of a less central hub, such as PG. The results of the present study emphasize the functional role of PG in the pathophysiology of SZ and highlight the advantage of its downregulation on mitochondrial function in SZ specimens, without detectable toxic effects *in-vitro*. Hence, PG may turn into a biomarker, and ultimately into a novel target for therapeutic intervention at least for a subgroup of SZ patients and possibly for additional disorders with impaired energy metabolism.

## Acknowledgment

The authors thank Dr. Maya Holdengreber and Dr. Ariel Shemesh, from Light Microscopy and Image Processing Unit of The Biomedical Core Facility at the Rappaport Faculty of Medicine, Technion, for their professional assistance, each in their respective fields.

## Author contribution statements

YL and DBS contributed to the designee, analysis and interpretation of the results, drafting and revising the paper. YL performed the imaging and MEA recordings, the latter performed with the guidance of OB. YL designed and constructed the plasmids, and with the help of RK performed the molecular work. DBS conceived the study, acquired funding and supervised the work.

